# APOE4 status is related to differences in memory-related brain function in asymptomatic older adults: Baseline analysis of the PREVENT-AD task fMRI dataset

**DOI:** 10.1101/2019.12.11.871681

**Authors:** Sheida Rabipour, Sricharana Rajagopal, Elsa Yu, Stamatoula Pasvanis, John Breitner, PREVENT-AD Research Group, M. Natasha Rajah

**Author notes:** Data used in preparation of this article were obtained from the Pre-symptomatic Evaluation of Novel or Experimental Treatments for Alzheimer’s Disease (PREVENT-AD) program (https://douglas.research.mcgill.ca/stop-ad-centre), data release 5.0 (November 30, 2017). A complete listing of PREVENT-AD Research Group can be found in the PREVENT-AD database: https://preventad.loris.ca/acknowledgements/acknowledgements.php?date=[2019-06-03]. The investigators of the PREVENT-AD program contributed to the design and implementation of PREVENT-AD and/or provided data but did not participate in analysis or writing of this report. Corresponding Author: M. Natasha Rajah, Ph.D., Room 2114, CIC Pavilion, Douglas Hospital Research Centre, Verdun, QC, H4H 1R3.

## Abstract

Episodic memory decline is one of the earliest symptoms of late-onset Alzheimer’s Disease (AD) and older adults with the apolipoprotein E e4 (+APOE4) genetic risk factor for AD may exhibit altered patterns of memory-related brain activity years prior to initial symptom onset. In the current study we report the baseline episodic memory task fMRI results from the PRe-symptomatic EValuation of Experimental or Novel Treatments for Alzheimer’s Disease (PREVENT-AD) study in Montreal, Canada, in which 327 healthy older adults, within 15 years of the parent’s conversion to AD, were scanned. During the task fMRI protocol volunteers were scanned as they encoded *and* retrieved object-location spatial source associations. The task was designed to discriminate between brain activity related to successful spatial source recollection and failures in spatial source recollection, with memory for only item (object) memory. Multivariate task-related partial least squares (task PLS) was used to test the hypothesis that +APOE4 adults with a family history of AD would exhibit altered patterns of brain activity in the recollection-related memory network, comprised of medial frontal, parietal and medial temporal cortices, compared to APOE4 non-carriers (-APOE4). We also tested for group differences in the correlation between event-related brain activity and memory performance in +APOE4 compared to -APOE4 adults using behavioral-PLS (B-PLS). We found group similarities in memory performance and in task-related brain activity in the recollection network. However, the B-PLS results indicated there were group differences in brain activity-behavior correlations in ventral occipito-temporal, medial temporal, and medial prefrontal cortices during episodic encoding. These findings are consistent with previous literature on the influence of APOE4 on brain activity and provide new perspective on potential gene-based differences in brain-behavior relationships in people with parental history of AD. Future research should further investigate the potential to distinguish risk of AD development based on memory performance and associated patterns of brain activity.

## 1. Introduction

Late-onset sporadic Alzheimer’s Disease (AD) accounts for an estimated 70% of dementia cases worldwide (Alzheimer’s Association, 2019; World Health Organization, 2019). Typically appearing after 65 years of age, this form of AD greatly increases in incidence beyond 85 years of age and has grown in prevalence with increasing life expectancy (Rabinovici, 2019). Despite the rising prevalence and burden of AD, there remain no widely effective treatments to prevent or delay symptom progression (Alzheimer’s Association, 2019). Moreover, new AD drug trials fail at a rate of over 100:1 (GBD Dementia Collaborators, 2019).

Given the current challenges of developing effective treatments for AD, increasing focus surrounds early identification, intervention, and prevention of AD in asymptomatic adults at higher risk of developing the disease (Frankish & Horton, 2017). Such risk factors include having an apolipoprotein E ε4 (APOE4) allele as well as first-degree family history (i.e., parent or sibling) of the disease (Donix, Small, & Bookheimer, 2012). Evidence further indicates that subtle neurological changes may precede symptom onset by as many as 20 years, during a period known as the ‘silent’ preclinical stage of AD (Brookmeyer, Abdalla, Kawas, & Corrada, 2018; Kern et al., 2018). Therefore, studies have increasingly focused on searching for early neural biomarkers or cognitive indices that could help predict AD development – particularly in individuals at higher risk of AD – at a period in which neurocognitive systems remain relatively intact (Leoutsakos, Gross, Jones, Albert, & Breitner, 2016; Molinuevo et al., 2016; Weiner & Veitch, 2015). Such early risk identification would permit access to interventions aimed at preventing or delaying AD onset (e.g., physical and cognitive exercise, control of blood pressure and cardiovascular conditions, improved diet, etc.) before the disease has progressed, and may help decrease the social and economic burdens associated with AD (Crous-Bou, Minguillon, Gramunt, & Molinuevo, 2017; National Academies of Sciences, 2017; Weimer & Sager, 2009).

Declines in memory of past personal events (i.e., episodic memory) represent one of the earliest symptoms of AD (Gardiner, 2001) and associate with preclinical or pre-symptomatic AD-related neuropathology (Bateman et al., 2012). Thus, the neural systems supporting episodic memory may be key candidate sites in which the first signs of AD-related neuropathology arise during the ‘silent’ phase of the disease (Backman, Small, & Fratiglioni, 2001; Sperling et al., 2010). Consistent with this theory, differences in task-related brain activity in people with mild cognitive impairment and AD, compared to healthy older adults, appear to reflect declines in memory and attention processing (Morrison, Rabipour, Knoefel, Sheppard, & Taler, 2018; Morrison, Rabipour, Taler, Sheppard, & Knoefel, 2019), and may index changes related to AD progression (Genon et al., 2013; Westerberg et al., 2013). Moreover, brain activity related to memory encoding in regions that subserve episodic memory – including hippocampal, parahippocampal, posterior parietal, and lateral prefrontal cortex – appears different in healthy aging, mild cognitive impairment, and AD (Zamboni et al., 2013). Notably, although recollection of rich contextual details related to a past event tends to decline even in healthy aging (Cansino et al., 2013; Spaniol & Grady, 2012), recognition of previously encountered objects or events appears more severely impacted in pathological aging (Wolk, Manning, Kliot, & Arnold, 2013). Episodic memory tasks that can differentiate the neural systems associated with familiarity, compared to recollection, may therefore be particularly helpful in identifying early signs of AD related neuropathology in healthy, at-risk adults.

Our past work demonstrated an association between episodic memory performance and changes in task-related fMRI detectable as early as middle-age (Ankudowich, Pasvanis, & Rajah, 2016) and more pronounced in APOE4 carriers or those with family history of AD (Rajah et al., 2017). These distinct associations appear in the absence of measurable behavioral differences and may be greater for the encoding and retrieval of contextual details associated with a past event, compared to general sense of familiarity (Ankudowich et al., 2016).

Examining brain-behavior relationships during episodic memory task performance may therefore provide new insights for identifying early AD indices. In particular, exploring differences in episodic memory-related brain function in older adults with and without risk factors such as an APOE4 allele and family history of AD may help better understand the impact of AD on memory systems (e.g., Meyer et al., 2018; Tardif et al., 2018; Villeneuve et al., 2018; Vogel et al., 2018).

### The present Study

Here we primarily aimed to report data from the functional neuroimaging task used in the PREVENT-AD program (https://douglas.research.mcgill.ca/stop-ad-centre). Moreover, whereas previous studies have largely focused on region-of-interest approaches with univariate analytical tools, comparatively few have examined the relationship between APOE4 and whole-brain functional changes. Thus, we further sought to examine the potential influence of carrying an APOE4 allele (i.e., +APOE4) on whole-brain activity during encoding and retrieval of objects (i.e., recognition) and their location (i.e., source recall), in cognitively healthy older adults with parental history of AD. We hypothesized that, relative to -APOE4 individuals, +APOE4 individuals would display different patterns of brain activity unrestricted to the hippocampus or medial temporal lobe during episodic encoding and retrieval. We further anticipated that the difference in task-related brain responses between +APOE4 vs. -APOE4 individuals would interact with object recognition and recall of spatial context. Specifically, given that source recollection performance and associated brain activity may change even in healthy aging (Cansino et al., 2013; Spaniol & Grady, 2012), we anticipated that differences in brain-behavior relationships would primarily reflect a deficit in object recognition in +APOE4 compared to - APOE4 individuals.

## 2. Methods

### 2.1 Participants: PREVENT-AD Cohort

Participants were recruited for the longitudinal PRe-symptomatic EValuation of Experimental or Novel Treatments for Alzheimer’s Disease (PREVENT-AD) program, an observational cohort study in Montreal, Canada (Breitner, Poirier, Etienne, & Leoutsakos, 2016). We evaluated baseline data from 327 older adults (*M*_age_=63.40 ± 5.24 years, 234 women) who were enrolled up to August 31^st^, 2017 (i.e., data release 5.0) and participated in the task fMRI portion of the study (see below).

We excluded participants on the basis of confounding genetic factors (i.e., APOE2 carriers, *n*=34; APOE44 homozygotes, *n*=7; unavailable genotype, *n*=3); having below-chance performance or fewer than eight trials per response type in the task fMRI protocol (*n*= 95); and poor fMRI image resolution (*n*=32). Our final sample comprised 172 older adults (*M*_age_=63.42 ± 5.12 years; 128 women), including 103 APOE33 homozygotes (i.e., -APOE4; 76/103=74% women) and 69 APOE34 heterozygotes (i.e., +APOE4; 52/69=75% women).

### 2.2 Protocol

Enrolment criteria for the PREVENT-AD trial are described elsewhere (Breitner et al., 2016). Briefly, all participants had at least one parent or multiple siblings diagnosed with sporadic AD or a condition suggesting Alzheimer’s-like dementia within 15 years (Tschanz et al., 2013). At baseline and during each subsequent follow-up assessment, participants performed neuropsychological tests as well as an object-location memory task in the scanner, described below. Here we focus on baseline analyses of the task-related fMRI based on APOE4 genotype. We describe longitudinal analyses of this task fMRI protocol in a forthcoming report. For more information on PREVENT-AD, see: douglas.qc.ca/page/prevent-alzheimer-the-centre.

### 2.3 Determination of Family History of AD

A brief questionnaire from the Cache County Study on Memory Health and Aging (Utah, USA) determined that all participants had a parent or multiple siblings: i) who had troubles with memory or concentration that was sufficiently severe to cause disability or loss of function; ii) for whom the condition had insidious onset or gradual progression and was not an obvious consequence of a stroke or other sudden insult.

### 2.4 APOE Genotyping

Genetic characterization was completed via blood draw, as previously described (Gosselin et al., 2016). DNA was isolated from 200 µl of the blood sample using QIASymphony and the DNA Blood Mini QIA kit (Qiagen, Valencia, CA, USA). APOE gene variant was determined using pyrosequencing with PyroMark Q96 (Qiagen, Toronto, ON, Canada).

### 2.5 Neuropsychological Testing

Neuropsychological assessments took approximately 40 minutes to administer and were completed prior to every testing session. Different versions of the RBANS were used in follow-up sessions to prevent practice effects. The test battery included:

*The Alzheimer-Dementia Eight Scale (AD8)*, an eight-item screening tool. The AD8 items index memory, orientation, judgment, and function. A score of two or above suggests impaired cognitive function (Galvin et al., 2005).
*The Clinical Dementia Rating (CDR)*, a five-point scale used to characterize memory, orientation, judgment & problem solving, community affairs, home & hobbies, and personal care (Berg, 1984). The information for each rating is obtained through a semi-structured interview of the patient and a reliable informant or collateral source (e.g., family member).
*The Montreal Cognitive Assessment (MoCA)*, a brief cognitive screening tool sensitive to mild declines in cognitive function (Nasreddine et al., 2004).
*The Repeatable Battery for the Assessment of Neuropsychological Status (RBANS)*, a battery of neuropsychological assessments aiming to identify abnormal cognitive decline in older adults (Randolph, Tierney, Mohr, & Chase, 1998). The RBANS provides scaled scores for five cognitive indices: immediate memory, visuospatial construct, language, attention, and delayed memory. We included these scaled scores, as well as the total score, in our analyses.

### 2.6 Task fMRI: Behavioral Protocol

Participants were instructed to lie supine in a 3T Siemens Trio scanner (see below), while performing a source memory task programmed in E-Prime version 1.0 (Psychology Software Tools, Inc; Figure 1). During an initial encoding phase, participants were cued (10s) to memorize a series of 48 colored line drawings of common objects from the BOSS database (Brodeur, Guerard, & Bouras, 2014) in their spatial location (i.e., to the left or right of a central fixation cross). Each object was presented for 2000ms followed by a variable inter-trial interval (ITI; durations of 2200, 4400, or 8800ms; mean ITI = 5.13s) to add jitter to the fMRI data collection (Dale & Buckner, 1997). Following the encoding phase, there was a 20-min delay during which participants received structural MRI scans.

**Fig. 1.**
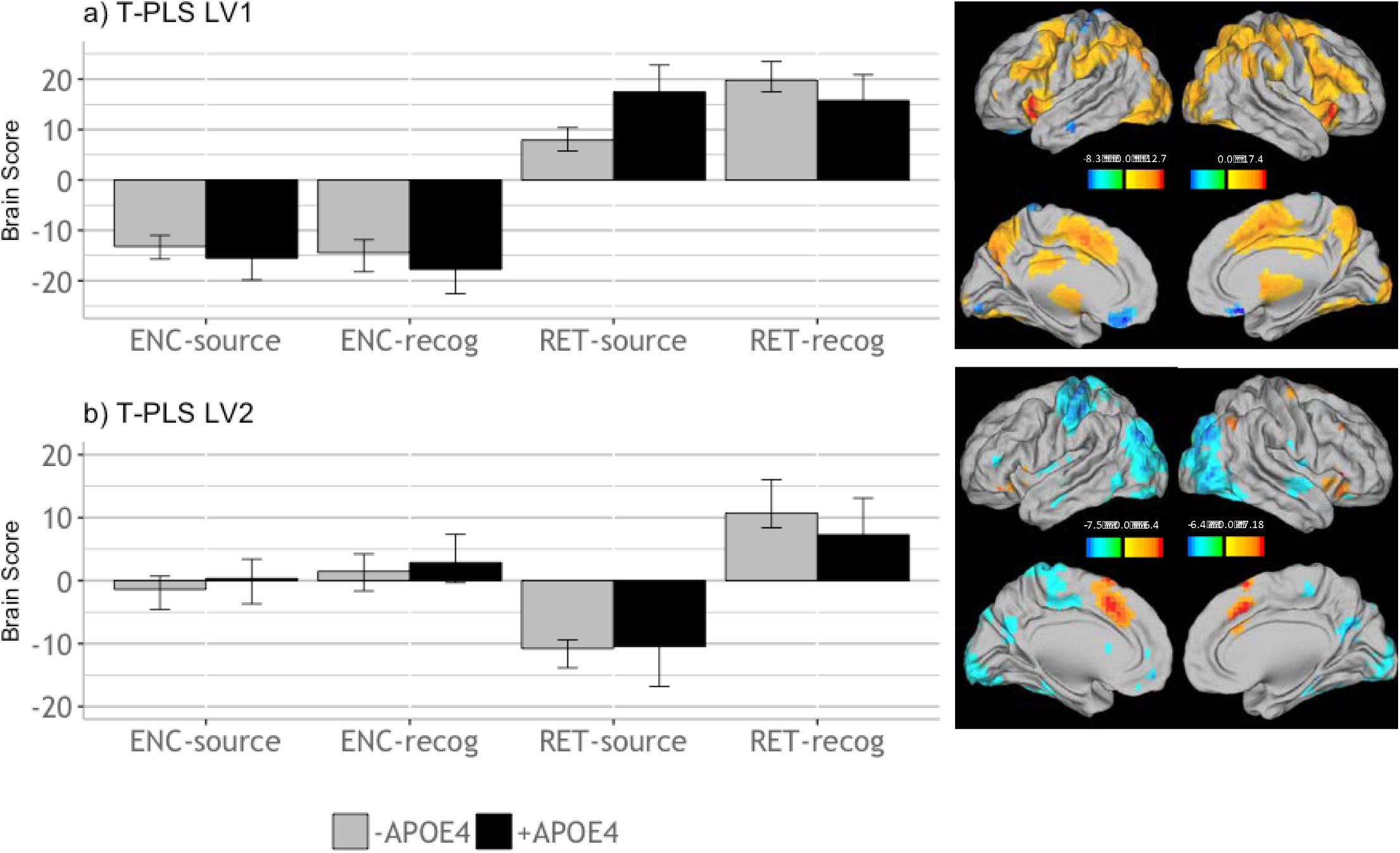
Design salience plot and singular image representing brain activity patterns by condition for (a) LV1 and (b) LV2, revealed by the T-PLS analysis. LEFT: Error bars on design salience plots represent 95% confidence intervals. RIGHT: Singular images were thresholded at a bootrstrap ratio of ±3.5, *p*<.001. Red brain regions represent positive brain saliences; blue regions represent negative brain saliences. Activations are presented on template images of the lateral and medial surfaces of the left and right hemispheres of the brain using Caret software (http://brainvis.wustl.edu/wiki/index.php/Caret:Download).

**Fig. 2.**
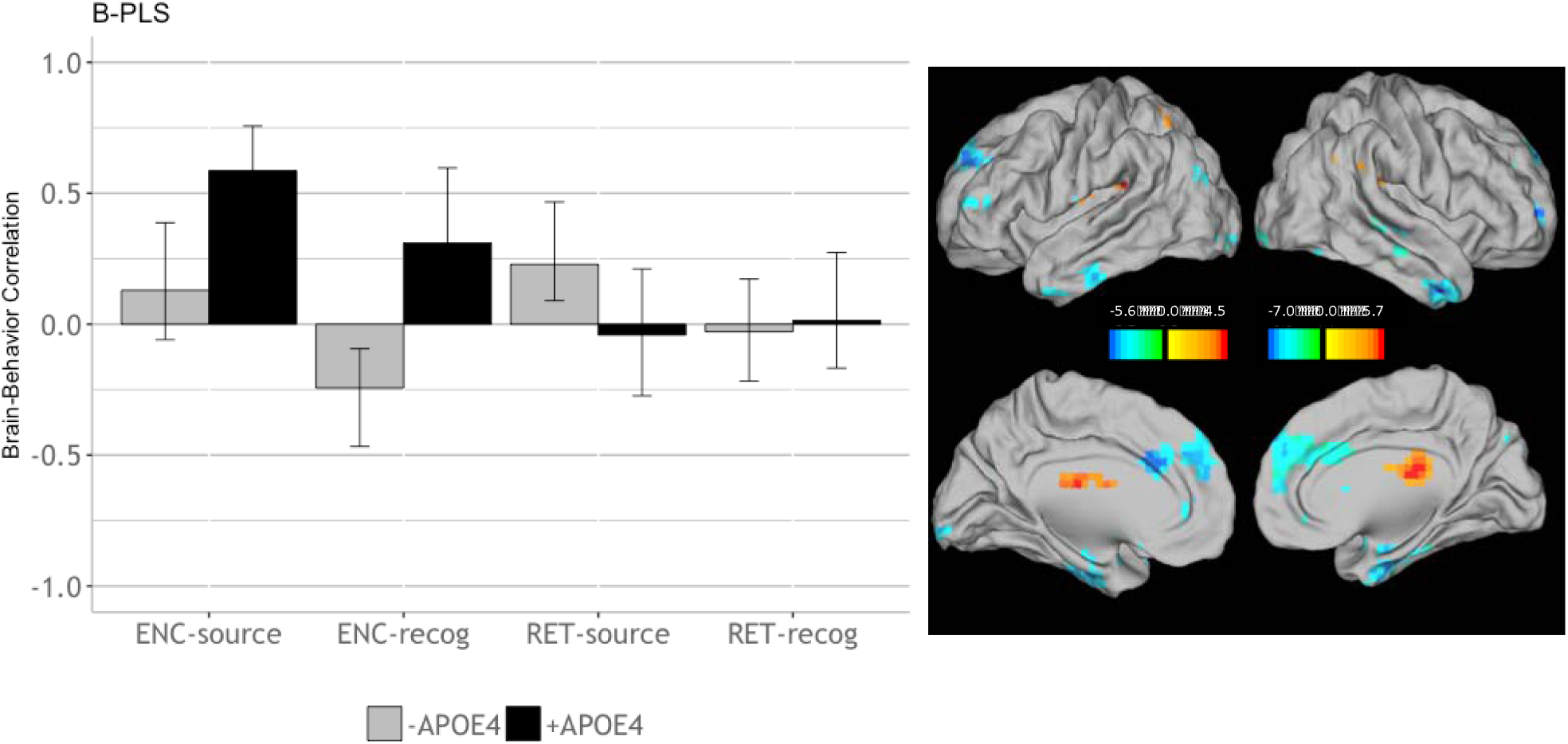
Correlations between brain activity and task performance by condition, revealed by the B-PLS analysis. LEFT: Bars represent brain-behavior correlations for each group, by condition. Error bars represent 95% confidence intervals. RIGHT: The singular image thresholded at a bootrstrap ratio of ±3.5, *p*<.001. Red brain regions represent positive brain saliences; blue regions represent negative brain saliences. Activations are presented on template images of the lateral and medial surfaces of the left and right hemispheres of the brain using Caret software (http://brainvis.wustl.edu/wiki/index.php/Caret:Download).

Following the 20-minute delay, a cue (10s) alerted participants to the beginning of the retrieval phase. During retrieval, participants were presented with 96 colored drawings of common objects: 48 ‘old’ (i.e., previously encoded) stimuli and 48 novel objects, in randomized order. Each object was presented in the center of the screen for 3000ms, with variable ITI (2200, 4400, or 8800ms). All participants used a fiber-optic 4-button response box to make task-related responses, and had an opportunity to familiarize themselves with the response choices during a practice session prior to testing. For each retrieval object, participants made a forced-choice between four-alternative answers: i) *“The object is FAMILIAR but you don’t remember the location”*; ii) *“You remember the object and it was previously on the LEFT”*; iii) *“You remember the object and it was previously on the RIGHT”*; and iv) *“The object is NEW”*. Thus (i) responses reflected object recognition, which may be more related to the utilization of familiarity vs. recollection based retrieval processes, (ii) and (iii) responses reflect associative recollection of object-location associations, and (iv) responses reflected either correct rejections of novel objects or failed retrieval (“misses”). Responding (i)-(iii) to new objects reflected false alarms.

### 2.7 fMRI data acquisition

Functional magnetic resonance images were acquired with a 3T Siemens Trio scanner using the standard 12-channel head coil, located at the Douglas Institute Brain Imaging Centre in Montreal, Canada. T1-weighted anatomical images were acquired after the encoding phase of the fMRI task using a 3D gradient echo MPRAGE sequence (TR=2300 msec, TE=2.98 msec, flip angle=9°, 176 1mm sagittal slices, 1×1×1 mm voxels, FOV=256 mm). Blood Oxygenated Level Dependent (BOLD) images were acquired using a single-shot T2* -weighted gradient echo-planar imaging (EPI) pulse sequence with TR=2000 msec, TE=30 msec, FOV=256 mm. Brain volumes with 32 oblique slices of 4mm thickness (with no slice gap) were acquired along the anterior-posterior commissural plane with in-plane resolution of 4×4 mm.

A mixed rapid event-related design was employed to collect task-related blood oxygen level dependent (BOLD) activation during performance of the memory task (see above). Visual task stimuli were generated on a computer and back-projected onto a screen in the scanner bore. The screen was visible to participants lying in the scanner via a mirror mounted within the standard head coil. Participants requiring correlation for visual acuity wore plastic corrective glasses.

### 2.8 Data Analysis

#### 2.8.1 Preprocessing of fMRI data

We converted reconstructed images to NIfTI format and preprocessed them using in Statistical Parametric Mapping software version 12 (SPM12). Images from the first 10s of scanning were discarded to allow equilibration of the magnetic field. All functional images were realigned to the first image and corrected for movement artifacts using a 6-parameter rigid body spatial transform and a partial least squares approach. Functional images were then spatially normalized to the MNI EPI-template using the “Old Normalize” method in SPM12 at 4×4×4 mm voxel resolution, and smoothed using an 8 mm full-width half-maximum (FWHM) isotropic Gaussian kernel. Participants with head motion exceeding 4mm in the x, y, or z axis during encoding and retrieval were excluded from further analyses. Participants with movements that could not be sufficiently repaired, resulting in distorted brain images as judged by an examiner, were excluded from further analysis. To be included in further analyses, all participants were required to have a minimum of eight observations per event type (i.e., object recognition and source recollection).

#### 2.8.2 Behavioral analyses

We performed behavioral data analyses on neuropsychological tests and episodic memory task performance using SPSS version 24 with a significance threshold of *p*=.05, Greenhouse-Geisser corrections for sphericity, and Bonferroni corrections for multiple comparisons, where applicable. Because evidence suggests meaningful sex/gender differences in the neural and behavioral correlates of episodic memory and cognition in aging (Gur & Gur, 2002; McCarrey, An, Kitner-Triolo, Ferrucci, & Resnick, 2016; Subramaniapillai et al., 2019), we also investigated self-reported sex as a factor in our analyses.

##### 2.8.2.1 Neuropsychological tests

We tested for group differences in AD8, MoCA, CDR, and RBANS scores using multivariate analysis of variance (MANOVA), with genotype (APOE34, APOE33) and self-reported sex (male, female) as independent factors, and test scores as dependent variables.

##### 2.8.2.2 Episodic memory task

We calculated mean accuracy and mean reaction time (RT; msec) for +APOE4 and - APOE4 participants, as well as self-reported males and females, for all possible response types: correct object recognition (recognizing old objects but providing no or incorrect associative spatial context), correct associative spatial context recollection (correctly recalling object-location associations); source misattributions (incorrectly identifying spatial context of an old object; e.g. saying an object previously seen on the left was initially presented on the right), false alarms (incorrectly identifying new objects as old), misses (incorrectly identifying old objects as new), and correct rejections (correctly identifying new objects). These stimulus-response categories are presented in Table 1. We computed overall accuracy (i.e., hits) using the sum of correct associative spatial context recollection judgments and correct object recognitions, including source misattribution trials. We used d’, computed as overall standardized hit rate minus standardized false alarm rate, as a measure of sensitivity, and c, computed by multiplying the average of the standardized hit and false alarm rates by -1, to measure response bias (Stanislaw & Todorov, 1999). We further estimated episodic memory performance by measuring the probability of source recall (pSource) and recognition (pRecog), calculated as follows:

1. pSource = Z_(source hits)_ – Z_(source misattributions)_
2. pRecog = Z_(recognition hits)_ – Z_(false alarms)_

**Table 1.** Participant responses and their categorization based on the presented stimuli.

| Stimulus | Participant Response |  |  |  |
| --- | --- | --- | --- | --- |
|  | <u>“Old, Left”</u> | <u>“Old, Right”</u> | <u>“Familiar”</u> | <u>“New”</u> |
| Old, Left | Source Hit | Source Misattribution | Recognition | Miss |
| Old, Right | Source Misattribution | Source Hit | Recognition | Miss |
| New | False Alarm | False Alarm | False Alarm | Correct Rejection |

Similar to d’, these scores provide a relative measure of accuracy that takes into account both hits and false alarms while isolating responses based on source recall versus familiarity. We then used multivariate general linear models to evaluate between-group differences on hits, pSource and pRecog scores, and response times (RT) on hit, miss, and false alarm trials.

#### 2.8.3 fMRI analyses

We used spatio-temporal partial least squares (PLS) to conduct multivariate event-related fMRI analysis using PLSGUI software (https://www.rotman-baycrest.on.ca/index.php?section=84). We selected PLS due to its ability to identify whole-brain spatially and temporally distributed patterns of brain activity that differ across experimental conditions and/or relate to a specific behavioral measure (McIntosh, Chau, & Protzner, 2004). We conducted two types of PLS analyses: i) mean-centered task partial least squares (T-PLS) to identify group similarities and differences in event-related activity during successful object-location associative encoding and retrieval; and ii) behavioral partial least squares (B-PLS) to examine group similarities and differences in the correlations between event-related activity and performance, indexed by pSource and pRecog. Details on PLS have been published elsewhere (Krishnan, Williams, McIntosh, & Abdi, 2011; McIntosh & Lobaugh, 2004).

For both T-PLS and B-PLS, we averaged the event-related data for each participant across the entire time series and stacked these data by group (i.e., APOE4) and based on participants’ subsequent memory performance as follows: i) encoding objects in which participants subsequently remembered object-location source associations (correct source recall; ENC-source); ii) encoding objects in which participants subsequently remembered only the object identified, but failed to recall spatial source information (source failure with object recognition; ENC-recog); iii) retrieval objects for which participants correctly recalled object-location source associations (RET-source); and iv) retrieval objects for which participants correctly recalled only the object identity, but failed to recall spatial source association (RET-recog). The stacked data matrix contained the fMRI data for each event onset (time lag = 0) with seven subsequent time lags, representing a total of 14s of activation after event onset (TR = 2s 7 = 14 s) for successfully *encoded* (ENC-source, ENC-recog) and successfully *retrieved* (RET-source and RET-recog) events. All participants analyzed had a minimum of eight correct events per event type. There was no signal at lag 0 because data were baseline corrected to the event onset. Therefore, signal in subsequent lags was expressed as percentage deviation from event onset.

##### 2.8.3.1 Task PLS (T-PLS)

We mean centered the fMRI data column-wise during the encoding and retrieval phases of the episodic memory task, to evaluate whole-brain similarities and differences between - APOE4 vs. +APOE4 individuals in brain activity related to encoding and retrieval of object-location associations. PLS performs singular value decomposition on the stacked data matrix to express the cross-covariance between the fMRI data and each condition into a set of mutually orthogonal latent variables (LVs). The number of LVs produced is equivalent to the number of event types included in the analysis. Thus, this analysis yielded eight LVs (4 event-types * 2 groups). Each LV comprises: i) a singular value reflecting the amount of covariance accounted for by the LV; ii) a design salience with a set of contrasts representing the relationship between tasks in each group and the pattern of brain activation; and iii) a singular image representing the numerical weights assigned to each voxel at each TR/time lag (i.e., the “brain salience”), yielding a spatio-temporal pattern of whole-brain activity for the entire time series. Design saliences and brain saliences can be either positive or negative: positive brain saliences are positively correlated to positive design saliences, whereas negative brain saliences are negatively correlated with positive design saliences (and vice-versa; Krishnan et al., 2011; McIntosh & Lobaugh, 2004). Thus, the pattern of whole brain activity identified by the singular image is symmetrically associated with the contrast effect identified by the design salience plot.

##### 2.8.3.2 Behavioral PLS

We used B-PLS to analyze whole-brain similarities and differences in brain activity directly correlated with pSource and pRecog during encoding and retrieval between +APOE4 and -APOE4 individuals. We stacked the behavioral vector containing pSource and pRecog in the same order as the fMRI data matrix (i.e., participant within group). As in the T-PLS, B-PLS performed singular value decomposition of the stacked cross-correlation matrix to yield eight LVs. Conversely, rather than design saliences, B-PLS analysis yields: i) a singular value, reflecting the amount of covariance explained by the LV; ii) a singular image consisting of positive and negative brain saliences, and iii) a correlation profile depicting how participants’ retrieval accuracy (pSource or pRecog) correlates with the pattern of brain activity identified in the singular image. The correlation profile and brain saliences represent a symmetrical pairing of brain-behavior correlation patterns for each group to a pattern of brain activity, respectively. As with the T-PLS analysis, brain saliences can have positive or negative values, and reflect whether activity in a given voxel is positively or negatively associated with the correlation profile depicted.

We assessed the significance of each LV in the T-PLS and B-PLS through 1000 permutations involving resampling without replacement from the data matrix to reassign the order of event types within participant. We determined the stability of the brain saliences using 500 bootstrap samples for the standard errors of voxel saliences for each LV, sampling participants with replacement while maintaining the order of event types for all participants. We ≥3.28 times greater than the bootstrap standard error (approximately corresponding to p=.001) and a minimum spatial extent of 10 contiguous voxels as stable.

We computed temporal brain scores for each significant LV to determine the time lags with the strongest correlation profile. Temporal brain scores reflect how strongly each participant’s data reflected the pattern of brain activity expressed in the singular image in relation to its paired correlation profile, at each time lag. We report only peak coordinates from time lags at which the correlation profile was maximally differentiated within the temporal window sampled (lags 2-5; 4-10s after event onset). We converted these peak coordinates to Talairach space using the icbm2tal transform (Lancaster et al., 2007) as implemented in GingerAle 2.3 (Eickhoff et al., 2009). Because our acquisition incompletely acquired the cerebellum, peak coordinates from this region are not reported. We used the Talairach and Tournoux atlas (Talairach & Tournoux, 1998) to identify the Brodmann area (BA) localizations of significant activations.

## 3. Results

After removing age outliers (i.e., individuals with age above or below 2SD from the mean), we analyzed data from 165 participants, including 98 -APOE4 and 67 +APOE4 individuals. Participant demographics and mean motion during fMRI are shown in Table 2. Evidence suggests that proximity to the age of AD diagnosis in a parent or sibling may predict the onset of dementia symptomatology (Villeneuve et al., 2018). We therefore included estimated years to AD symptom onset (EYO), calculated as the age of AD onset in the earliest affected family member subtracted from the participant’s current age, as part of these baseline analyses.

**Table 2.** Demographic background and mean fMRI motion (in mm) of participants included in behavioral analyses (n=165) based on APOE4 group, represented as mean values *±* standard deviation.

|  | Years of Age | n Women:Men | Years of Education | EYO | Motion |
| --- | --- | --- | --- | --- | --- |
| -APOE4 | 63.43 ± 4.51* | 73 : 25 | 14.80 ± 3.34 | 9.93 ± 8.08 | .025 ± .067 |
| +APOE4 | 61.98 ± 3.91 | 51 : 16 | 15.48 ± 3.46 | 9.19 ± 7.66 | .027 ± .066 |

\* $p=.033$

We found that -APOE4 individuals were significantly older (t_163_=2.15, *p*=.033).We therefore included age as a covariate in our analyses of performance below. Moreover, although groups were balanced in their sex distributions (*X^2^*=.057, *p*=.812), we found a significant correlation between sex and performance on the MoCA (*r*_pb_ = .227, *p* = .003) and CDR (*r*_pb_ = -.183, *p* = .019). Thus, we included sex as a factor in the corresponding analyses, We found no other significant effects of age or sex at baseline.

### 3.1 Neuropsychological Performance

We found a significant effect of APOE4 (F_(1,159)_=10.07, *p*=.002, η_p_^2^ =.06) and of sex (F_(1,159)_=6.45, *p*=.011, η_p_^2^ =.04) on MoCA total score, a significant effect of APOE4 on AD8 total score (F_(1,160)_=6.24, *p*=.014, η_p_^2^ =.038; ns after correcting for multiple comparisons), and a significant effect of sex on CDR total score (F_(1,160)_=5.21, *p*=.024, η_p_ =.032; ns after correcting for multiple comparisons). Follow up analyses revealed that +APOE4 participants had significantly higher MoCA scores compared to -APOE4 individuals (t_162_=2.81, *p*=.006), and that women had significantly higher MoCA scores compared to men (t_58.11_=2.66, *p*=.01). Notably, nine -APOE4 individuals scored below the clinical cutoff of 26 compared to only two +APOE4 individuals, although this difference was not significant (*X*^2^=1.31, *p*=.25). Conversely, the proportion of women who scored below 26 on the MoCA (5/119=4%) was significantly lower than the proportion of men (7/34=21%; *X*^2^=7.77, *p*=.005). Our analyses revealed no significant differences in RBANS test scores between APOE4 groups (Wilk’s λ =.971, F(6,148) =.744, *p*=.615) or based on sex (Wilk’s λ =.981, F(6,148) =.487, *p*=.817) at baseline. These results are presented in Table 3.

**Table 3.** Neuropsychological performance, represented as mean values ± standard deviation. Data missing from one participant who did not receive the MoCA (n=164) and seven participants who did not receive the RBANS (n=158).

|  | AD8 | CDR | MoCA | Immediate Memory | Visuospatial Construction | Language | Attention | Delayed Memory | RBANS Total |
| --- | --- | --- | --- | --- | --- | --- | --- | --- | --- |
| -APOE4 | .12 ± .41 | .015 ± .09 | 27.68**<br>± 1.62 | 104.34<br>± 10.9 | 94.76<br>± 12.3 | 98.40 ± 10.19 | 106.03 ± 15.74 | 104.41 ± 8.73 | 101.62 ± 9.36 |
| +APOE4 | .24 ± .61 | .007 ± .06 | 28.38 ± 1.45 | 103.73 ± 11.23 | 94.66<br>± 13.12) | 101.66 ± 8.69 | 105.41 ± 13.78 | 103.47 ± 10.95 | 102.05 ± 10.73 |

\*\* $p=.006$

**Table 4.**
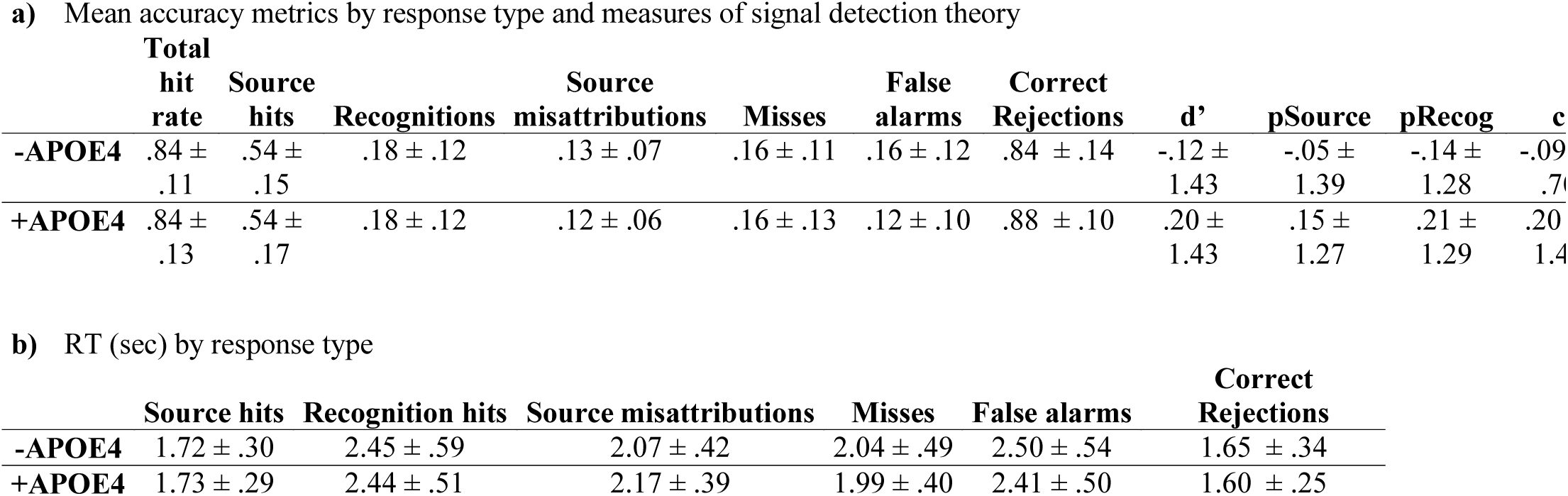
Performance on the episodic memory task (n=165). Errors are standard deviations.

|  | **Total**<br>**hit**<br>**rate** | **Source**<br>**hits** | **Recognitions** | **Source**<br>**misattributions** | **Misses** | **False**<br>**alarms** | **Correct**<br>**Rejections** | **d'** | **pSource** | **pRecog** | **c** |
| --- | --- | --- | --- | --- | --- | --- | --- | --- | --- | --- | --- |
| **-APOE4** | .84 ±<br>.11 | .54 ±<br>.15 | .18 ± .12 | .13 ± .07 | .16 ± .11 | .16 ± .12 | .84 ± .14 | -.12 ±<br>1.43 | -.05 ±<br>1.39 | -.14 ±<br>1.28 | -.09<br>.70 |
| **+APOE4** | .84 ±<br>.13 | .54 ±<br>.17 | .18 ± .12 | .12 ± .06 | .16 ± .13 | .12 ± .10 | .88 ± .10 | .20 ±<br>1.43 | .15 ±<br>1.27 | .21 ±<br>1.29 | .20<br>1.4 |

b) RT (sec) by response type
|  | **Source hits** | **Recognition hits** | **Source misattributions** | **Misses** | **False alarms** | **Correct**<br>**Rejections** |
| --- | --- | --- | --- | --- | --- | --- |
| **-APOE4** | 1.72 ± .30 | 2.45 ± .59 | 2.07 ± .42 | 2.04 ± .49 | 2.50 ± .54 | 1.65 ± .34 |
| **+APOE4** | 1.73 ± .29 | 2.44 ± .51 | 2.17 ± .39 | 1.99 ± .40 | 2.41 ± .50 | 1.60 ± .25 |

### 3.2 fMRI: Episodic Memory Task Performance

Behavioral results from the object-location associative memory task are shown in 4. We found no significant group differences in task performance (Wilk’s λ=.969, F_(5,156)_ =.983, *p*=.43) or RT (Wilk’s λ=.972, F_(7,147)_ =.607, *p*=.749) at baseline. Our analyses nevertheless revealed significant differences in RT based on response type (F_(3.99,610.8)_ =3.37, *p*=.01, η_p_^2^=.022). Participants had significantly longer RT for recognition (i.e., source failure) and FA trials compared to source hits, source misattributions, misses, and correct rejections (t_164_≥6.45, *p*≤.0001), and significantly faster RT for correct rejections compared to all other trials (t_164_≥4.00, *p*≤.0001). Of trials presenting “old” (i.e., previously viewed) objects, participants had the fastest RT for source hits (t_159_≥8.98, *p*≤.0001).

### 3.3 fMRI results

#### 3.3.1 T-PLS

The PLS analysis yielded two significant LVs (*p*<.0001). The first significant LV accounted for 66.7% of the cross-block covariance and identified brain regions in which activity significantly differed during encoding, compared to retrieval, in both +APOE4 and -APOE4 groups (Figure 1A). Table 5 lists the local maxima from LV 1. In both APOE4 groups, positive salience brain regions were more active during retrieval, compared to encoding, whereas the negative salience brain regions were more active during encoding, compared to retrieval, across all groups. More activity was detected in the left ventromedial/orbitofrontal/anterior cingulate, and the left lateral middle temporal cortex during encoding, compared to retrieval. Conversely, during retrieval there was more activity in the bilateral claustrum, cingulate gyrus, inferior frontal gyrus, and precuneus, as well as the left middle frontal gyrus, compared to encoding.

**Table 5.**
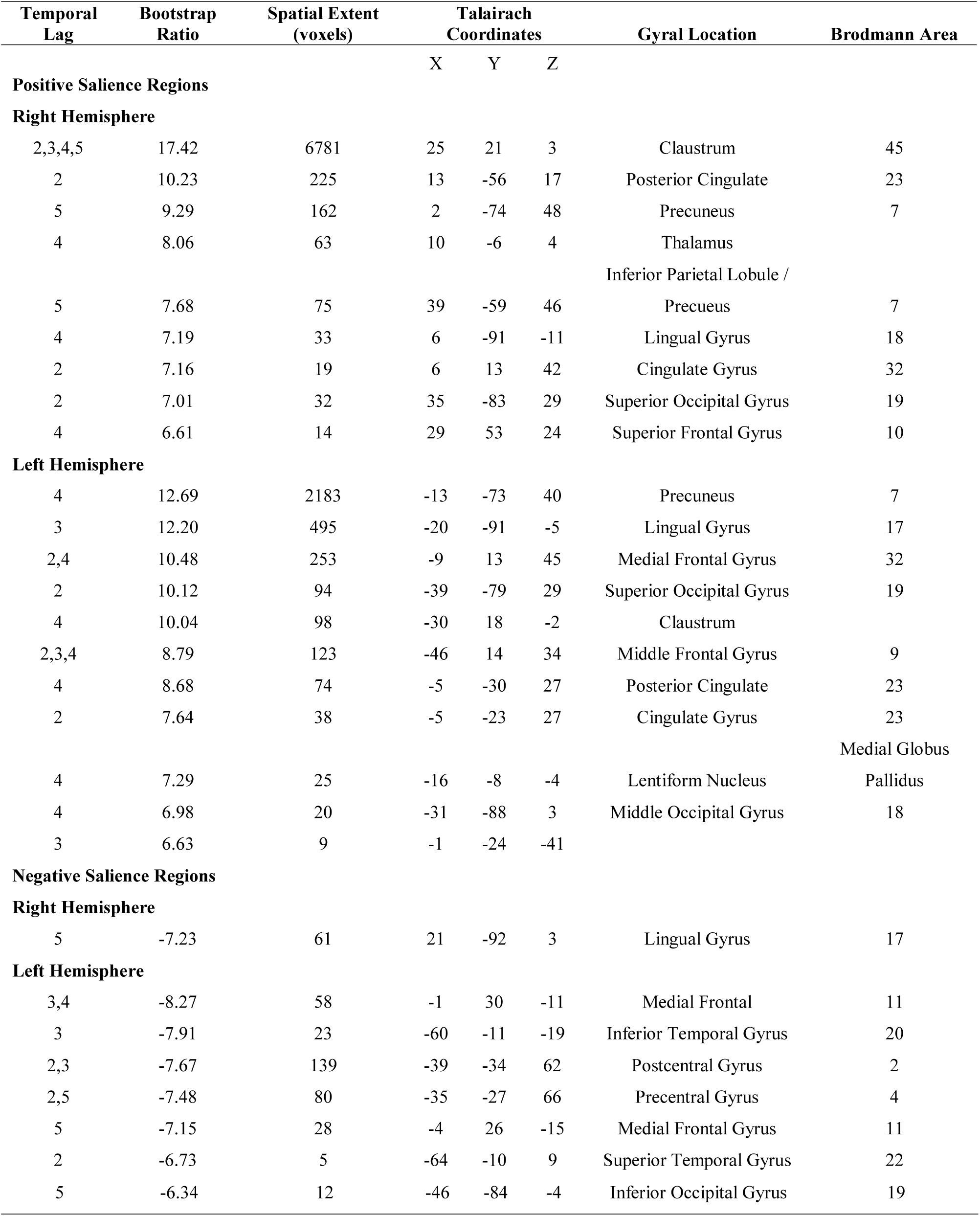
Local maxima revealed for LV1 of the T-PLS analysis. We report only lags 2-5, and clusters with a spatial threshold of at least 10 continuous voxels.

LV2 of the T-PLS accounted for 13.78% of the cross-block covariance. The design salience plot and singular image presented in Figure 1B indicates that this LV identified brain regions that were differentially activated during correct retrieval of object-location associations, compared to objects alone (i.e., source hits vs. source misattributions and failures) in both APOE groups. The local maxima from this LV are presented in Table 6. This LV identifies mainly negative brain saliences and indicates that activity was greater in middle occipital, inferior parietal lobule, caudate, and temporal gyri, including the left parahippocampus and right hippocampus, during object-location associative retrieval, compared to object only retrieval. In contrast, activity was greater in the bilateral inferior frontal gyrus, cingulate gyrus, left middle frontal gyrus, and right inferior parietal lobule during object only retrieval, compared to object-location retrieval.

**Table 6.** Local maxima revealed for LV2 of the T-PLS analysis. We report only lags 2-5, and clusters with a spatial threshold of at least 10 continuous voxels.

| Temporal Lag | Bootstrap Ratio | Spatial Extent (voxels) | Talairach Coordinates |  |  | Gyral Location | Brodmann Area |
| --- | --- | --- | --- | --- | --- | --- | --- |
|  |  |  | X | Y | Z |  |  |
| Positive Salience Regions |  |  |  |  |  |  |  |
| Right Hemisphere |  |  |  |  |  |  |  |
| 4,5 | 7.18 | 306 | 2 | 17 | 42 | Cingulate Gyrus | 32 |
| 4, 5 | 5.87 | 127 | 40 | 21 | 0 | Inferior Frontal Gyrus | 47 |
|  |  |  |  |  |  | Inferior Parietal |  |
| 5 | 4.75 | 39 | 46 | -62 | 42 | Lobule | 40 |
| 4,5 | 4.55 | 32 | 32 | -15 | 61 | Precentral Gyrus | 6 |
| Left Hemisphere |  |  |  |  |  |  |  |
| 3,4 | 6.39 | 160 | -45 | 17 | 2 | Inferior Frontal Gyrus | 47 |
| 5 | 4.84 | 113 | -45 | 33 | -8 | Inferior Frontal Gyrus | 47 |
| Negative Salience Regions |  |  |  |  |  |  |  |
| Right Hemisphere |  |  |  |  |  |  |  |
| 3,4,5 | -6.37 | 127 | 14 | 23 | 18 | Caudate |  |
| 2 | -5.96 | 192 | 28 | -77 | 37 | Precuneus | 19 |
|  |  |  |  |  |  | Superior Temporal |  |
| 2,3,4 | -5.37 | 82 | 51 | -8 | -6 | Gyrus | 22 |
|  |  |  |  |  |  | Inferior Temporal |  |
| 2 | -5.40 | 332 | 43 | -69 | -2 | Gyrus | 37 |
| 2 | -4.86 | 100 | 47 | -3 | 12 | Insula | 13 |
| 3 | -4.79 | 35 | 54 | -10 | 44 | Postcentral Gyrus | 3 |
| 4 | -4.78 | 24 | 32 | -35 | -6 | Hippocampus |  |
|  |  |  |  |  |  | Parahippocampal |  |
| 2,5 | -4.05 | 53 | 29 | -30 | -20 | Gyrus | 36,37 |
| 2 | -3.91 | 19 | 13 | -12 | 36 | Cingulate Gyrus | 24 |
| Left Hemisphere |  |  |  |  |  |  |  |
| 2,3,4 | -7.54 | 953 | -35 | -30 | 62 | Precentral Gyrus | 4 |
|  |  |  |  |  |  | Middle Occipital |  |
| 3 | -7.34 | 2294 | -31 | -79 | 22 | Gyrus | 19 |
|  |  |  |  |  |  | Inferior Parietal |  |
| 2 | -6.62 | 1747 | -39 | -32 | 47 | Lobule | 40 |
| 3,5 | -6.23 | 175 | -20 | 24 | 13 | Caudate |  |
| 4 | -5.99 | 493 | -23 | 32 | 3 |  |  |
| 3 | -5.43 | 70 | -60 | -12 | -5 | Middle Temporal | 21 |
|  |  |  |  |  |  | Gyrus |  |
| 5 | -5.03 | 74 | 21 | -44 | 15 |  | 5 |
| 5 | -4.85 | 35 | -17 | -42 | 65 | Postcentral Gyrus | 5 |
|  |  |  |  |  |  | Parahippocampal |  |
| 2,4,5 | -4.79 | 29 | -38 | -34 | -10 | Gyrus | 36 |
| 2 | -4.50 | 95 | 29 | 32 | 4 |  |  |
| 2 | -4.32 | 16 | -16 | 43 | 12 | Medial Frontal Gyrus | 10 |
|  |  |  |  |  |  | Inferior Temporal |  |
| 2 | -4.31 | 11 | -56 | -7 | -15 | Gyrus | 20 |
| 2,5 | -4.07 | 10 | -19 | -6 | -29 | Uncus | 36 |
| 5 | -4.01 | 10 | -31 | -73 | 0 | Lingual Gyrus | 18 |
| 2 | -3.94 | 22 | -12 | -21 | 13 | Thalamus |  |

#### 3.3.2 B-PLS Results

After removing performance outliers based on pSource and pRecog, we analyzed data from 129 participants (78 -APOE4 and 51 +APOE4). The B-PLS yielded one significant LV (*p*=.008), which accounted for 23.82% of the cross-block covariance (Figure 4). Table 7 lists the local maxima for positive and negative salience brain regions. This LV identified brain regions with group differences in the correlation between pRecog performance and encoding activity for objects which participants subsequently retrieved only object identity but failed to recall the spatial source information (ENC-recog). In addition, this LV identified brain regions in which activity during object-location source retrieval (RET-source) correlated with pSource performance in -APOE4 adults. Specifically, in -APOE4 individuals, activity in positive salience brain regions during ENC-recog events was negatively correlated with pRecog scores, and activity in these same regions during RET-source events was positively correlated with pSource scores. In addition, in -APOE4 adults, activity in negative salience brain regions during ENC-recog events was positively correlated with pRecog scores and activity in these regions during RET-source was negatively correlated with pSource scores. Positive salience brain regions included right thalamus, right supramarginal gyrus and left insula (Table 7). Negative salience brain regions included bilateral occipito-temporal cortices, uncus, caudate, and anterior-medial PFC, as well as left parahippocampal gyrus. Therefore, in -APOE4 individuals activity in a traditional episodic memory network (negative salience brain regions) during encoding predicted subsequent object-only retrieval (i.e., source failures), and less activation in these regions at retrieval predicted better source performance.

**Table 7.** Local maxima revealed by the B-PLS analysis. We report only lags 2-5, and clusters with a spatial threshold of at least 10 continuous voxels

| Temporal Lag | Bootstrap Ratio | Spatial Extent<br>(voxels) | Talairach Coordinates |  |  | Gyral Location | Brodmann Area |
| --- | --- | --- | --- | --- | --- | --- | --- |
|  |  |  | X | Y | Z |  |  |
| Positive Salience Regions |  |  |  |  |  |  |  |
| Right Hemisphere |  |  |  |  |  |  |  |
| 3, 5 | 5.69 | 63 | 58 | -53 | 29 | Supramarginal Gyrus | 39,40 |
| 3 | 5.46 | 319 | 6 | -26 | 20 | Thalamus |  |
| Left Hemisphere |  |  |  |  |  |  |  |
| 3 | 4.50 | 17 | -38 | -33 | 19 | Insula | 13 |
| Negative Salience Regions |  |  |  |  |  |  |  |
| Right Hemisphere |  |  |  |  |  |  |  |
| 2,3,5 | -7.02 | 442 | 6 | 48 | 38 | Medial Frontal Gyrus | 6,8,10 |
| 2 | -5.28 | 42 | 44 | 5 | -27 | Middle Temporal Gyrus | 21 |
| 3,4,5 | -5.16 | 155 | 22 | -2 | -28 | Uncus | 28,36, Amygdala |
| 3 | -4.96 | 47 | 14 | 17 | 6 | Caudate | Caudate Body |
| 5 | -4.84 | 12 | 43 | -31 | -2 | Superior Temporal Gyrus | 22 |
| 5 | -4.22 | 34 | 21 | -87 | -11 | Middle Occipital Gyrus | 18 |
| Left Hemisphere |  |  |  |  |  |  |  |
| 3 | -5.61 | 50 | -16 | 13 | 16 | Caudate | Caudate Body |
| 3,5 | -5.44 | 143 | -34 | -14 | -27 | Uncus | 20,28 |
| 2 | -4.86 | 23 | -20 | 25 | 35 | Middle Frontal Gyrus | 8 |
| 5 | -4.71 | 18 | -53 | -18 | -24 | Fusiform Gyrus | 20 |
| 3 | -4.39 | 14 | -9 | -90 | -12 | Lingual Gyrus | 18 |
| 3 | -4.35 | 64 | -23 | 34 | 22 | Medial Frontal Gyrus | 9 |
| 3 | -4.33 | 16 | -2 | 15 | 67 | Superior Frontal Gyrus | 6 |
| 3 | -3.97 | 11 | -41 | 13 | -31 | Superior Temporal Gyrus | 38 |
| 3 | -3.89 | 13 | 43 | -39 | -2 |  |  |
| 2 | -3.78 | 10 | -27 | -46 | -1 | Parahippocampal Gyrus | 19 |

Adults with +APOE4 genotype exhibited the opposite pattern of brain activity-behavior correlations to -APOE4 adults at encoding, and did not exhibit significant brain-behavior correlations at retrieval. Specifically, in +APOE4 adults, activity within right thalamus, right supramarginal gyrus and left insula during ENC-recog and ENC-source correlated with better subsequent memory for both event types. In contrast, encoding activity in more traditional episodic memory-related areas (negative salience regions) was negatively correlated with subsequent memory in these adults.

Taken together these results indicate that +APOE4 and -APOE4 adults engaged distinct patterns of brain activity at encoding to support subsequent memory. In addition, -APOE4 adults appeared to exhibit an encoding/retrieval flip in the regions supporting memory.

#### 3.3.3 Post-hoc regions of interest (ROI) analyses

We investigated activation profiles for the ROI displaying the highest peaks during encoding in our B-PLS analysis and consistent with our TPLS results (Fig. 3). Multivariate general linear models evaluating group differences in ROI activation, averaged across lags 2-5, indicated no significant group differences during ENC-source or ENC-recog.

**Fig. 3.**
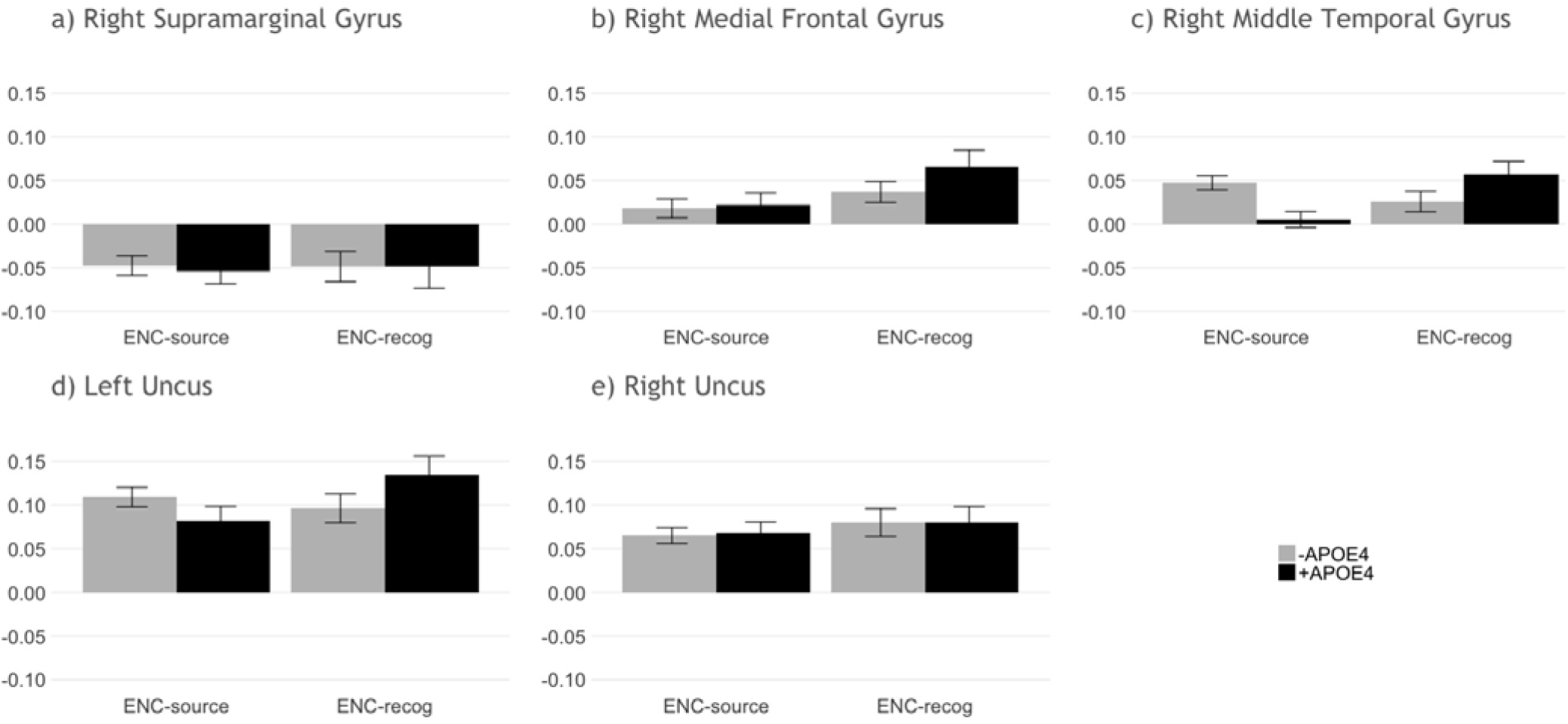
Mean activation across lags 2-5 at encoding for ROI identified via B-PLS. Bars represent standard error.

## 4. Discussion

The PREVENT-AD study aimed to identify signs of AD pathogenesis in pre-symptomatic older adults with elevated risk of AD (Breitner et al., 2016). Because neural changes may occur decades before the onset of clinical AD symptoms (Brookmeyer et al., 2018; Kern et al., 2018), PREVENT-AD used sensitive biomarkers such as brain imaging, biochemistry, metabolic, and cognitive measures to explore characteristic changes during disease progression. The PREVENT-AD study directly addressed several shortcomings of previous investigations of prodromal AD (O’Donoghue, Murphy, Zamboni, Nobre, & Mackay, 2018), including use of sufficiently large sample size, appropriate sample composition (e.g., older adults with family history of AD), longitudinal design, and comprehensive cognitive and biological outcomes.

Here we drew on baseline data from the PREVENT-AD cohort to examine the potential influence of having an APOE4 allele on episodic memory function. Few studies have assessed the relationship between APOE4 and brain activity from the whole-brain perspective. Instead studies have often used univariate approaches and focused on specific regions of interest (i.e., the medial temporal lobe system) that are implicated in AD pathology (e.g., Bassett et al., 2006; Dennis et al., 2010; Filippini et al., 2009; Johnson et al., 2006; Xu et al., 2009). These studies have contributed significantly to our understanding of memory-related dysfunction in preclinical samples with the APOE4 risk factor for AD. The current study aimed to expand on this extant literature by using a data-driven multivariate approach, PLS, to compare brain activity related to episodic memory encoding and retrieval between -APOE4 and +APOE4 individuals with family history of AD. In addition, contrary to previous studies focusing primarily on recognition/novelty, we employed a novel episodic memory task distinguishing object-location source association from recognition via a single response.

### 4.1.1 Few Behavioral Group Differences

Behaviorally, we found that both -APOE4 and +APOE4 individuals performed well on neuropsychological tests as well as our episodic memory task, with greater MoCA scores in +APOE4 individuals. These results largely align with previous studies showing no significant behavioral differences in cognitive performance on the basis of APOE4 status in younger (Taylor et al., 2017) and healthy older adults of similar demographic background (Reas et al., 2019), as well as our previous work using a similar episodic memory task in middle-aged adults at risk of AD (Rajah et al., 2017). However, given the high sensitivity of the MoCA for detecting mild cognitive impairment (Pinto et al., 2019), lower MoCA performance among - APOE4 participants may support theories suggesting that carrying an APOE4 allele may paradoxically benefit cognitive function in early and midlife (Evans et al., 2014). The majority of participants in both groups scored well above the cutoff of 26, argued by some to be overly conservative even in highly educated older adults (Elkana, Tal, Oren, Soffer, & Ash, 2019), suggesting that most participants retained healthy cognitive performance. In addition, evidence suggests a greater effect of APOE4 in men compared to women (Qiu, Kivipelto, Aguero-Torres, Winblad, & Fratiglioni, 2004); this effect may have been masked in our largely female population. Moreover, both +APOE4 and -APOE4 participants were relatively young, with median ages of 61 years and 62, respectively, and were an estimated 9-10 years in proximity to symptom onset. This stage may therefore have been too early to detect detrimental changes in cognitive performance on the basis of APOE4 (Rawle et al., 2018).

Given the lack of clinical symptoms in our sample, we expected comparable behavioral performance in both groups. As predicted, we observed no significant effect of +APOE4 status on episodic memory performance. Across both groups, participants were more likely to recollect the object-location association (54%) than recognize the object only (i.e., fail to remember the source; 18%). Nevertheless, we observed a non-significant trend whereby - APOE4 participants had a greater number of FA and fewer correct rejections, as well as low response discriminability (i.e., negative d’, pSource, and pRecog values) and a bias towards selecting “old” (i.e., negative c value); this was not true of +APOE4 participants. Importantly, our task demanded that participants reflect on their memory of a presented object (or lack thereof) and respond in a single four-choice step. Therefore, during each retrieval trial where participants recognized an object as “old”, participants likely attempted to first recall the object-location source association, selecting that the object was simply familiar only if they failed to recall its contextual information. Supporting this interpretation, participants had longer RT during both trials where participants correctly recognized objects but failed to recall the source and where participants misidentified new objects as old (i.e., FA). Conversely, successfully encoded object-location source associations, as well as correct rejections, were identified much more rapidly. These behavioral patterns suggest that recognition trials in the present task reflect objects for which participants could not retrieve source information (i.e., source failures; see also our discussion of task activation and brain-behavior correlations below), with slightly better discriminability in +APOE4 individuals.

### 4.1.2 Few Group Differences in Task Activation Patterns

Results from our mean-centered T-PLS analysis support the effectiveness of our task in selectively isolating episodic memory encoding vs. retrieval (LV1) and object recognition vs. source recollection (LV2). Furthermore, counter to our hypothesis, T-PLS revealed group similarities in task-related activation during episodic encoding and retrieval, as well as for object recognition vs. source recollection.

Specifically, T-PLS LV1 demonstrated that both -APOE4 and +APOE4 older adults relied preferentially on activity in the ventromedial frontal and temporal cortices for encoding objects, and in the frontal and parietal cortices to retrieve objects from memory. These results align with previous reports of increased activation in left-lateralized medial frontal, middle and inferior temporal, as well as primary and secondary sensory cortices (e.g., prefrontal and ventro-occipito-temporal areas) for encoding objects and their location, compared to increased activation in broad bilateral areas including the precuneus, parietal, and medial prefrontal cortices during episodic retrieval in older adults (Maillet & Rajah, 2014; Salami, Eriksson, & Nyberg, 2012). Such prefrontal activation at encoding, coupled with decreased task-positive activation and task-negative deactivation, appears to correlate with successful recall in older adults (Maillet & Rajah, 2014), and may reflect greater strategic elaboration of episodic memories in older age (Shing, Brehmer, Heekeren, Backman, & Lindenberger, 2016).

Similarly, T-PLS LV2 indicated that, in both APOE4 groups, object-location source association correlated with broad activity in the bilateral caudate, precuneus, and temporal cortices, as well as left parahippocampal gyrus and right hippocampus. As with patterns identified in T-PLS LV1, these areas align with previous reports of increased activity in regions related to stimulus perception and medial temporal lobes – particularly in the left hemisphere – during successful encoding of contextual information (Maillet & Rajah, 2014), and overlap with areas of the “core recollection network” (Rugg & Vilberg, 2013). Notably, some studies have suggested a greater association between parahippocampal function and familiarity, rather than recollection; however, in contrast to our task, the tasks used in such reports have largely relied on subjective indices, including subjective judgments, sometimes coupled with confidence ratings (Daselaar, Fleck, & Cabeza, 2006). Conversely, we found that object recognition correlated with activity in the bilateral middle and medial frontal areas, and the right cingulate, angular gyrus, and inferior parietal lobule in both -APOE4 and +APOE4 individuals.

Our T-PLS results, coupled with our behavioral findings, suggest an alternative interpretation of recognition trials in the context of our task. On one hand, -APOE4 and +APOE4 individuals exhibited similar processing of object recognition and source recollection in areas previously shown to associate with episodic memory in aging (Ankudowich et al., 2016; Mitchell & Johnson, 2009). However, regions active during object recognition in the present study typically correlate with recollection of objects and their associated contextual information. The reversed pattern observed here may reflect the nature of our task design. Because the task required participants to select a single response indexing their memory of a presented object, trials in which participants indicated recognition without object-location associations may have reflected their failure to recall contextual information. This process may have led to activation of regions involved in recollection despite failed source recall. Our behavioral results support this interpretation: longer RT during both recognition and FA trials suggests that participants searched for object-location source association but, in both cases, failed to retrieve the correct information. Conversely, successfully encoded object-location source associations were retrieved much more rapidly (see discussion of behavioral results above).

Drawing on objective measures of source recall and recognition, our results extend previous findings, which have been largely identified using the traditional “remember vs. know” paradigm that often relies on subjective report and may be less sensitive to the effects of age on episodic memory (Koen & Yonelinas, 2014). Our results further indicate partial indices of pathological memory-related brain activity in older adults at higher risk of AD: on the one hand, similar to healthy older adults, participants in the present study demonstrated activation of parahippocampal cortices during encoding of novel stimuli (Zamboni et al., 2013). However, similar to high performing individuals with early dementia (Grady et al., 2003), our participants demonstrated greater prefrontal and temporal activation during episodic encoding and broad activation in areas affected by AD, including the claustrum, precuneus, inferior parietal, and middle and medial frontal cortices, during retrieval, suggesting possible preclinical indices of pathological aging.

Contrary to our hypothesis, our findings related to performance and task-related activation (i.e., T-PLS) support studies suggesting no measurable distinction in episodic memory performance and underlying brain function based on APOE4 status. Although APOE4 is linked with impaired memory, cognition, and functional activity, more rapid cognitive decline, and hippocampal atrophy in clinical populations (e.g., mild cognitive impairment and AD) and preclinical states such as subjective memory decline (Cosentino et al., 2008; Farlow et al., 2004; Striepens et al., 2011), effects in asymptomatic older adults are less clear. Studies have shown an association between APOE4 and accelerated memory decline later in life (Rawle et al., 2018) as well as altered structure and function in medial temporal lobe circuitry (Gallagher & Koh, 2011; Matura et al., 2014). Moreover, using verbal episodic memory tasks, previous studies have shown greater activation of areas affected in AD (e.g., dorsolateral prefrontal cortex and superior temporal gyrus) during episodic retrieval in young adults with at higher genetic risk of AD (Braskie et al., 2013). However, although APOE4 is the strongest genetic risk factor for late-onset AD (Hersi et al., 2017), presence of the allele does not appear to be a significant independent predictor of AD (Jessen et al., 2011; O’Donoghue et al., 2018). Moreover, lifestyle choices (e.g., higher education level, moderate alcohol consumption) may mitigate APOE4 effects and contribute to discrepant reports of its influence on cognitive aging (Reas et al., 2019). Although an investigation of lifestyle factors was beyond the scope of this study, high education among both APOE4 groups, coupled with our participants’ eagerness to engage in stimulating activities such as PREVENT-AD, may at least partly explain our lack of observed group differences in behavior and task-related fMRI.

### 4.1.3 APOE4-Based Differences in Brain-Behavior Correlations

In contrast to our performance and T-PLS results, and supporting our hypothesis, our B-PLS analysis indicated group differences in correlations between brain activity at encoding and subsequent memory performance. These differences were most pronounced for subsequently recognized objects, where successful recognition performance correlated with encoding-related activity in bilateral medial and middle frontal cortices, temporal areas, uncus, and caudate, as well as left parahippocampus in -APOE4 individuals, compared to right inferior frontal, superior temporal, and supramarginal cortices, as well as the left precuneus, claustrum, and insula in +APOE4 individuals. We also found differential reliance on brain activity supporting retrieval of object-location source associations in APOE4 carriers vs. non-carriers. Whereas performance correlated negatively with activity in regions typically associated with source recollection (e.g., bilateral occipito-temporal cortices, uncus, caudate, and anterior-medial PFC, as well as left parahippocampal gyrus in -APEO4 participants, this correlation was positive in +APOE4 individuals. Notably, we found few group differences in correlations between performance and brain activity during encoding of source memory (ENC-source), or recognition of objects in the absence of source recall (RET-recog).

These results support our hypothesis that group differences in brain activity would emerge for object recognition and suggest that source recall vs. recognition performance may associate with different neural substrates in -APOE4 vs. +APOE4 individuals. These patterns may reflect a difference in approach taken by -APOE4 and +APOE4 individuals to complete the task. For example, mnemonic strategy training appears to improve performance on face-name associative memory tasks and further correlate with increased frontoparietal activity in older adults with amnesic mild cognitive impairment (Simon et al., 2018, 2019). However, because task instructions did not direct participants to use any particular approach or strategy, and based on the lack of behavioral differences, this seems unlikely to explain the observed findings. In particular, the observed group differences in brain-behavior correlations align with the antagonistic pleiotropy hypothesis of APOE, which posits that APOE4 may benefit cognition early in life but associate with cognitive decline later in life, possibly mediated by frontal-executive processes (Han & Bondi, 2008; Tuminello & Han, 2011). Decreased performance-related medial temporal lobe activity coupled with increased activity in right frontal regions in +APOE4 participants, in the absence of behavioral deficits, may index compensatory neural recruitment to support over-burdened task-related areas. Such activity may reflect early signs of AD conversion in +APOE4 older adults. Further support for this theory would require longitudinal analyses of task-related brain-behavior correlations.

### 4.2 Limitations and Future Directions

Despite the relatively large number of older adults who participated in the PREVENT-AD study, we excluded many from the present analyses based on genetic composition, performance, and quality of MRI scans. Thus, different patterns may have emerged with a larger sample. However, our results are largely consistent with the previous literature on cognitive performance and task-related brain activity in -APOE4 vs. +APOE4 individuals. In addition, here we focused on task-related activity on successful trials because of an insufficient number of unsuccessful trials (i.e., false alarms and misses). Analyzing such trials, when possible, may provide greater insights into the mechanisms underlying changes in episodic memory function with elevated age and AD risk, as has previously been shown for recognition memory in healthy aging (Stevens, Hasher, Chiew, & Grady, 2008). Such analyses should represent a greater focus in future research. Similarly, the present study aimed to examine brain activation patterns associated with a single encoding and retrieval phase, carried out over a relatively short interval. However, using a comparable paradigm, researchers have identified that activity in distinct areas, including posteromedial areas at encoding and frontal areas at retrieval, supports durable memories that last over the course of weeks (Vidal-Pineiro et al., 2017). Such time course likely reflects a more representative period over which memories form and consolidate over the long term, and may serve as important regions of interest in future investigations, particularly of pathological aging. Finally, the present results reflect only the baseline data from a longitudinal cohort; our future analyses will integrate these findings with performance and memory-related brain activity from follow-up assessments.

## 5. Conclusions

Here we found different correlations between brain activity and subsequent memory performance in -APOE4 compared to +APOE4 individuals, despite similar neuropsychological and episodic memory performance as well as task-related activation during episodic encoding and retrieval for both recognized objects and recalled object-location associations. Consistent with our hypothesis, this difference in brain-behavior relationships was particularly prominent for recognition, compared to source memory. Together, our results suggest that -APOE4 and +APOE4 individuals recruit brain regions expected to be active during encoding and retrieval of object recognition and object-location associations in older adults; however, APOE4 status may influence the way in which brain regions that subserve episodic memory encoding support recognition performance. Moreover, our activation-related findings, coupled with performance outcomes, highlight the importance of task considerations in interpreting processes underlying source recall and suggest that recognition may represent a failure of source retrieval.

Our findings are consistent with our previous work on the influence of APOE4 on brain activity in middle-aged adults at risk of AD (Rajah et al., 2017) and are the first to identify recognition-related differences in brain-behavior relationships in asymptomatic older adults at risk of AD. These findings add a nuanced perspective to the study of neural processes associated with recollection vs. familiarity as well as the role of APOE4 on brain-behavior relationships in asymptomatic older adults with family history of AD, which remains poorly characterized (O’Donoghue et al., 2018). Future research should further investigate the lifespan effects of APOE4 on neurocognitive processes and the potential to distinguish risk of AD development based on genetic composure, behavior, and associated patterns of brain activity.

## Acknowledgments

We thank members of the Rajah lab, the Cerebral Imaging Centre, and the StoP-AD Centre at the Douglas Hospital Research Centre for their helpful feedback on this work. We also acknowledge the Fonds de Recherche Québec – Santé and Canadian Institutes of Health Research for their support of our work.

PREVENT-AD was launched in 2011 as a $13.5 million, 7-year public-private partnership using funds provided by McGill University, the Fonds de Recherche du Québec – Santé, an unrestricted research grant from Pfizer Canada, the Levesque Foundation, the Douglas Hospital Research Centre and Foundation, the Government of Canada, and the Canada Fund for Innovation. Private sector contributions are facilitated by the Development Office of the McGill University Faculty of Medicine and by the Douglas Hospital Research Centre Foundation (http://www.douglas.qc.ca/).

